# Coronacept – a potent immunoadhesin against SARS-CoV-2

**DOI:** 10.1101/2020.08.12.247940

**Authors:** Hadas Cohen-Dvashi, Jonathan Weinstein, Michael Katz, Maayan Eilon, Yuval Mor, Amir Shimon, Romano Strobelt, Maya Shemesh, Sarel J Fleishman, Ron Diskin

## Abstract

Angiotensin-converting enzyme 2 (ACE2) is the cellular receptor for severe acute respiratory syndrome coronavirus 2 (SARS-CoV-2). Computational analysis of mammalian ACE2 orthologues suggests various residues at the interface with the viral receptor binding domain that could facilitate tighter interaction compared to the human-ACE2. Introducing several mutations to the human-ACE2 resulted with significantly augmented affinity to the viral spike complex. This modified human-ACE2 fused to an Fc portion of an antibody makes a potent immunoadhesin that effectively targets SARS-CoV-2.

## Introduction

Coronavirus disease 2019 (COVID-19), caused by the severe acute respiratory syndrome coronavirus 2 (SARS-CoV-2), is an ongoing devastating pandemic leading to a substantial global death toll and an unprecedented economic loss. There is an urgent need for anti-viral countermeasures that will help to save lives as well as shorten the recovery and hospitalization time of sick people. Immunotherapy has the potential to neutralize viruses directly as well as to recruit immune effector functions to clear infected cells, and it makes a promising approach that was already shown to be beneficial in combating other viral diseases. A monoclonal antibody (mAb) against the respiratory syncytial virus is effective and was approved for treating infected children ^1,2^. A breakthrough in the field of HIV was the isolation of broadly-neutralizing mAbs ^3,4^ and the affirmation of their use in treating ^5,6^ and protecting ^7,8^ individuals from HIV-1. In addition, antibodies against Lassa ^9^, Junin ^10^, Ebola ^11^, and even SARS ^12^ viruses have been proven effective in animal models. Preliminary reports indicate that also in the case of COVID-19, sera from convalescent patients help to fight the disease^13,14^. Hence, targeted immunotherapy may become a powerful anti-viral tool for fighting COVID-19.

Though anti-SARS-CoV-2 neutralizing mAbs were recently isolated from convalescent patients ^15-17^, it would be important to diversify our immunotherapeutic toolkit to maximize the chances for success. Diversification may be achieved using immunoadhesins, which are antibody-like molecules that consist of a binding domain fused to an Fc portion on an antibody ^18^. The viral cellular receptor could serve as a binding domain for constructing such immunoadhesins. Due to natural adaptation, however, zoonotic viruses may bind to their animal-derived ortholog cellular receptors at higher affinities than the human cell-surface receptors ^19^. Thus, immunoadhesins that are constructed with the host-ortholog receptors can provide superior anti-viral therapeutics. We recently demonstrated this approach by constructing Arenacept, which is a powerful immunoadhesin that targets viruses from the *Arenaviridae* family of viruses ^20^.

SARS-CoV-2 is a zoonotic virus that utilizes angiotensin-converting enzyme 2 (ACE2) as a cellular receptor ^21-23^. The genome of SARS-CoV-2 is similar to bat-derived SARS-like coronaviruses ^24^, but the exact origin of the virus is unknown and other animal species may have served as intermediate reservoirs before SARS-CoV-2 crossed over to the human population ^25^. It was shown that ACE2 orthologs from various animals can serve as entry receptors for SARS-CoV-2 ^26^. Furthermore, it was demonstrated that the human-ACE2 is a suboptimal receptor for SARS-CoV-2 ^27^. We therefore reasoned that the human-ACE2 could be modified to make it a superior binder of SARS-CoV-2. Such an engineered ACE2 may be better suited for therapeutic applications than the human ACE2.

## Results

The binding of SARS-CoV-2 to its ACE2 receptors is mediated by the receptor-binding domain (RBD), which is part of its spike complex ^23,28-30^. ACE2 has a long helical segment at its N′ terminus, which forms most of the RBD-recognition site on ACE2 (Fig. 1a). A multiple sequence alignment of over 200 ACE2 sequences derived from mammals indicates that many of the ACE2 positions that comprise the SARS-CoV-2 recognition site are not conserved (Fig. 1b). This notion indicates an enormous putative sequence space that ACE2 can adopt.

**Fig. 1.**
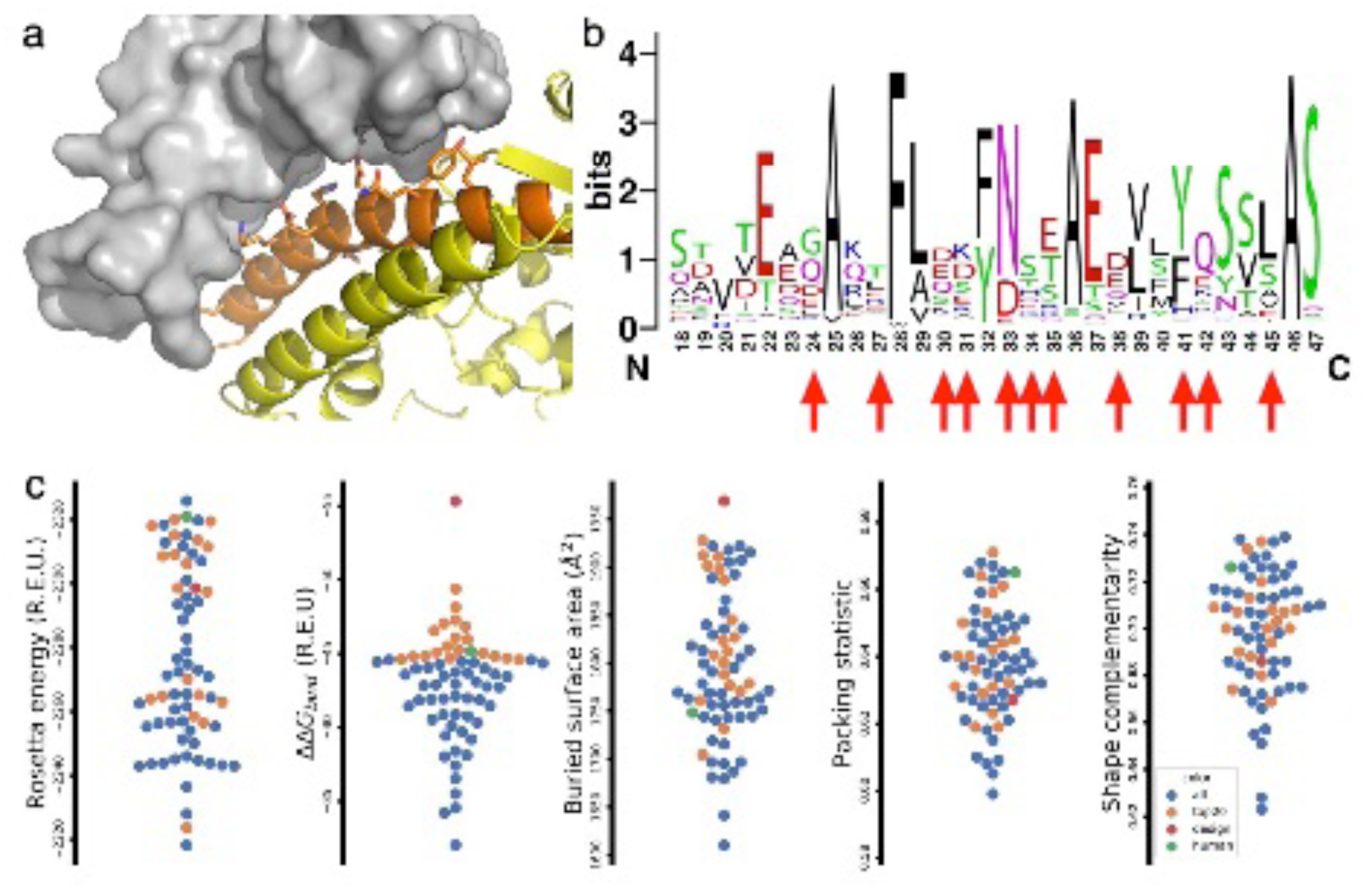
Diversity of the SARS-CoV-2/ACE2 interface. **a**. Structure of the SARS-CoV-2 RBD (grey surface) in complex with human-ACE2 (orange and yellow ribbon) (PDB: 6M17). The side chains of residues that make the recognition site for SARS-CoV-2 are shown as sticks. The N’-terminal helix of ACE2 that makes the central part of the binding site is highlighted in orange. **b**. The sequence diversity of the first N’-terminal helix of ACE2 in mammals is presented using a WebLogo display ^37^. The abundance of the amino acid types in each position is represented by the height of their single-letter code. The residues that interact with the RBD of SARS-CoV-2 are indicated by red arrows. **c**. Interface properties of the various RBS/ACE2-ortholog models. Each dot represents a single model. From left to right, five panels show the calculated total Rosetta energy (using Rosetta energy units), the binding energy (ΔΔ*G* for binding), the buried surface area, the packing statistics, and the shape complementarity of the interface. All the panels are arranged such that the values at the top represent better results. The RBD/human-ACE is indicated with a green dot. The modified ACE2 is indicated with a red dot.

To identify advantageous alterations of ACE2 that may enhance the binding to SARS-CoV-2, we selected 70 orthologous ACE2 genes with sequence identity to the human-ACE2 greater than 80%. We used Rosetta atomistic modelling calculations to assess the stability, binding energy (ΔΔ*G*_*bind*_), interface packing and shape complementarity for the RBD (starting from PDB entry 6VW1) (Fig. 1c) ^30^. We visually inspected the top-20 models according to ΔΔ*G*_*bind*_ and identified mutations that the calculations indicated would improve contacts with the SARS-CoV-2 RBD relative to the human ACE2. Due to the high sequence diversity in ACE2, many design options were available. We rejected mutations to Trp, due to their tendency to form undesired promiscuous interactions and furthermore consulted a recent deep mutational sequencing dataset on ACE2 mutations and their impact on binding to the RBD^27^ eliminating mutations that were depleted in this dataset.

Three of the mutations that we decided to incorporate are located at the first N’-terminal helix of human-ACE2. These three mutations include a T27L mutation that improves packing with hydrophobic residues of SARS-CoV-2 RBD (Fig. 2a), a D30E mutation that forms a new salt-bridge with Lys417 of SARS-CoV-2 RBD (Fig. 2b), and a Q42R mutation that may form a salt-bridge with Asp38 of ACE2 and stabilize it in a configuration that favors the formation of a hydrogen bond with Tyr449 of SARS-CoV-2 RBD (Fig. 2c). Alternatively, the new arginine may assume a different rotamer that makes favorable electrostatic interactions with the main-chain carboxylic oxygen of Gly447 from the SARS-CoV-2 RBD (Fig. 2c). Besides these three mutations at the N-terminal helix of ACE2 that we selected, we identified two additional sites in the surrounding regions of ACE2. In the first site, we identified a putative change of Glu75 and Leu79 to arginine and tyrosine, respectively, that may interact favorably with Phe486 of SARS-CoV-2 RBD (Fig. 2d). In the second site outside the first helix of ACE2, N330F may improve packing against the aliphatic portion of Thr500 from SARS-CoV-2 RBD (Fig. 2e). We used Rosetta to model the combination of these six mutations and its impact on the binding to SARS-CoV-2 RBD. Our design showed a remarkable improvement in ΔΔ*G* of binding as well as in the buried surface area (Fig. 1c).

**Fig. 2.**
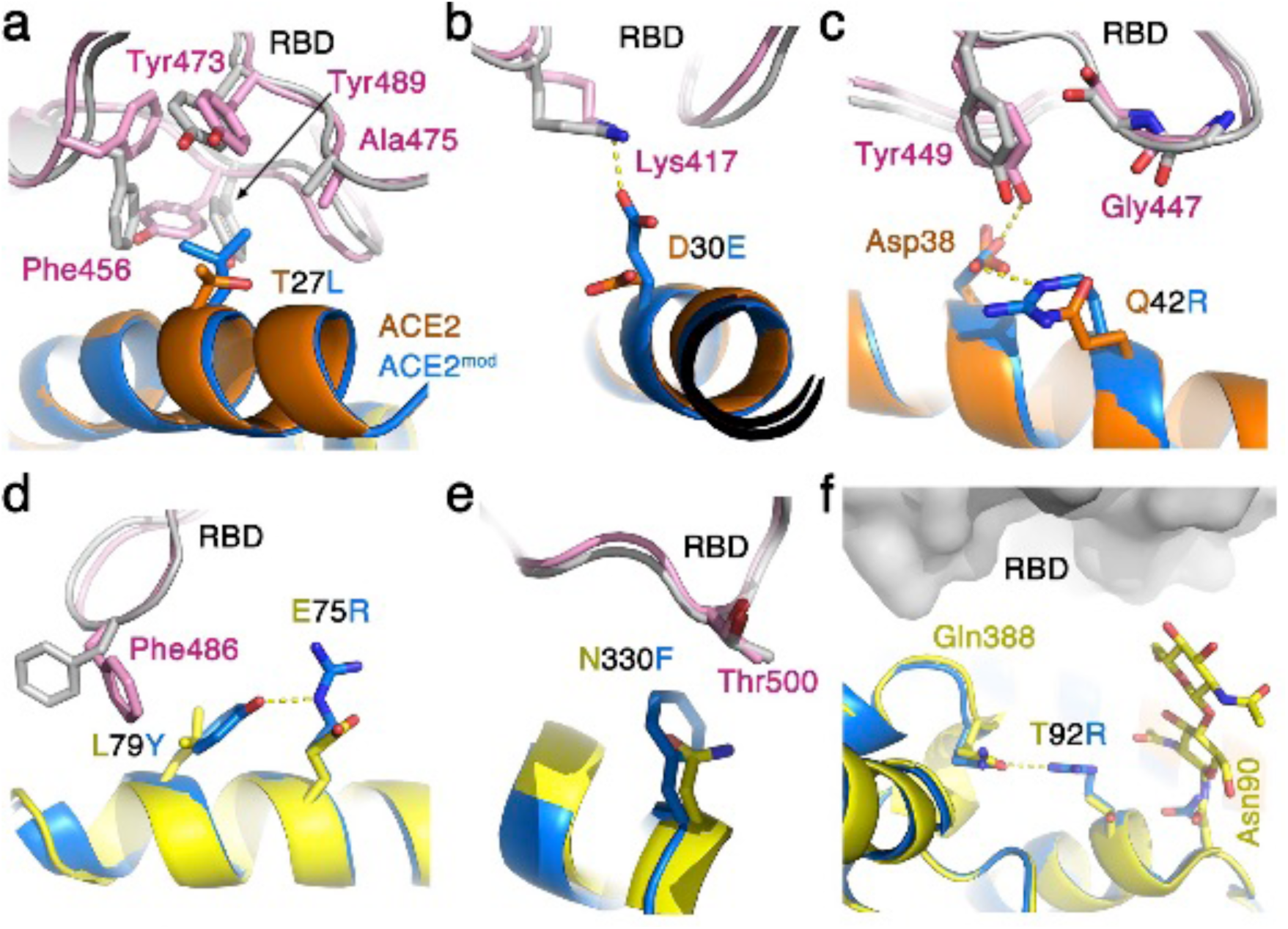
Optimized ACE2 interface for improved binding of SARS-CoV-2. The interfaces of SARS-CoV-2 RBD/human-ACE2 (grey and orange/yellow, respectively) and of the SARS-CoV-2 RBD/modified-ACE2 (pink and light blue, respectively) are shown. **a**. Leucine, instead of threonine in position 27, makes better Van der Waals interactions with hydrophobic residues on the SARS-CoV-2 RBD. **b**. Glutamic acid in position 30 of ACE2 can make a salt bridge with Lys417 of SARS-CoV-2, but not an aspartic acid that is present in the human-ACE2. **c**. Arginine in position 42 can form a salt-bridge with Asp38 of ACE2 to stabilize it in a configuration that allows it to make a hydrogen bond with the hydroxyl of Tyr449 from SARS-CoV-2 RBD. An arginine in this position can also assume a different rotamer that will allow it to form electrostatic interaction with the main-chain carbonyl oxygen of Gly447 of SARS-CoV-2 RBD. **d**. A double replacement of Leu79 and Glu75 with tyrosine and arginine respectively allows favorable interaction between Phe486 of SARS-CoV-2 RBD and Tyr79 that is stabilized through a hydrogen bond by Arg75. **e**. Phenylalanine in position 330 of ACE2 is predicted to pack better against the aliphatic portion of Thr500 from SARS-CoV-2 RBD. **f**. A replacement of Thr92 with arginine abrogates the glycosylation site on Asn90, which bears a glycan that can sterically interfere with the binding of SARS-CoV-2 RBD. An arginine in position 92 of ACE2 can form a hydrogen bond with the nearby Gln388.

We decided to incorporate additional modification at other sites on top of modifying ACE2 residues that directly interact with SARS-CoV-2 RBD. Human-ACE2 has a putative glycosylation site at Asn90 that was shown to bear a glycan according to the SARS-CoV-2 RBD/human-ACE2 cryo-EM structure ^21^. This N-linked glycan projects toward the SARS-CoV-2 RBD, and presumably imposes steric constraints for the binding of SARS-CoV-2 RBD. The aforementioned deep mutational scanning dataset^27^ is highly enriched with mutations in this N-linked glycosylation site, further supporting this notion. To eliminate this glycosylation site, we mutated Thr92 from the N-X-T glycosylation motif to an arginine that can make polar interactions with nearby glutamine (Fig. 2f). Besides serving as a cellular receptor for SARS-CoV-2, ACE2 is an enzyme that has a critical biological function in regulating blood pressure by hydrolyzing angiotensin II ^31^. Since the enzymatic activity of the engineered ACE2 may complicate its therapeutic use, we also mutated its key catalytic position Glu375 to leucine.

To test our design, we produced two chimeric proteins that included amino acids 19-615 of the human-ACE2 ectodomain (omitting the original signal peptide) fused to an Fc portion of human IgG1, with or without the eight above-mentioned mutations (i.e., T27L, D30E, Q42R, E75R, L79Y, N330F, T92R, & E375L). Both the WT construct (ACE2-Fc) and our designed ACE2 construct (ACE2^mod^-Fc) readily expressed as secreted proteins using HEK293F cells in suspension and were easily purified to near homogeneity using protein-A affinity chromatography (Fig. 3a). Testing the enzymatic activity of both ACE2-Fc and ACE2^mod^-Fc verified that the latter is indeed catalytic dead (Supplementary Fig. 1). We then immobilized the two immunoadhesins on a surface plasmon resonance sensor chip and used purified SARS-CoV-2 RBD as an analyte to determine their binding affinities, which is a configuration that does not allow avidity. A simple 1:1 binding model gave a good fit for the binding data of ACE2^mod^-Fc to SARS-CoV-2 RBD, but the binding of ACE2-Fc to SARS-CoV-2 RBD could not be fitted using this model, and we, therefore, used a more complex heterogeneous-ligand model that assumes some heterogeneity of the ACE2-Fc (Fig. 3b). Such heterogeneity could presumably originate from partial glycosylation at Asn90 of ACE2. Remarkably, the binding affinity of ACE2^mod^-Fc to SARS-CoV-2 RBD is more than two orders of magnitude stronger compared with the binding affinity of ACE2-Fc (Fig. 3b). While the association rate (*k*_a_) of ACE2^mod^-Fc to SARS-CoV-2 RBD is somewhat faster compared to ACE2-Fc, the improvement mainly stems from a dramatic two-orders of magnitude slower dissociation rate (*k*_d_).

**Fig. 3.**
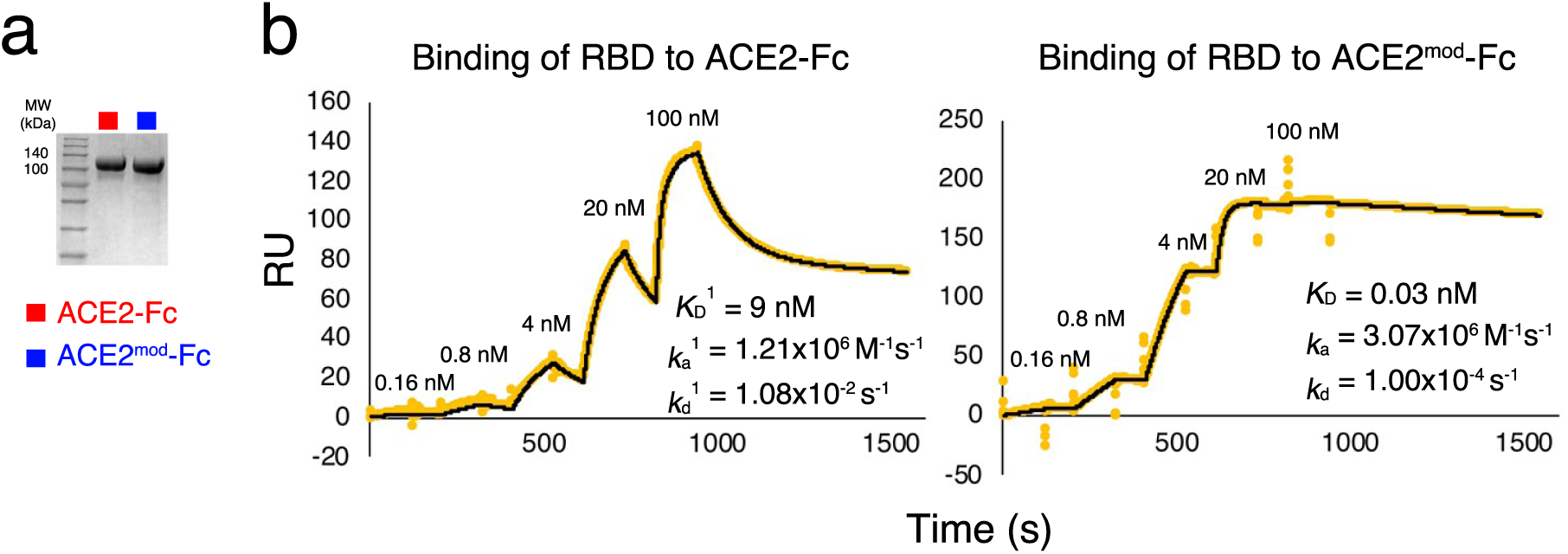
ACE2^mod^-Fc is a superior binder of SARS-CoV-2. **a**. Coomassie-stained SDS-PAGE showing ACE2-Fc and ACE2^mod^-Fc. **b**. SPR analyses of SARS-CoV-2 RBD interaction with ACE2-Fc and ACE2^mod^-Fc. Both ACE2-Fc and ACE2^mod^-Fc were immobilized to a protein-A sensor chip and SARS-CoV-2 RBD was injected at the indicated concentration series in a single-cycle kinetics experiment. The kinetic parameters are indicated. In the case of SARS-CoV-2 RBD binding to ACE2^mod^-Fc these parameters are derived from a simple 1:1 binding model. In the case of SARS-CoV-2 RBD binding to ACE2-Fc, these parameters are derived from heterogeneous-ligand binding model, and reflect the first component. These experiments were repeated twice and a representative sensorgram is shown for each ligand.

To test if the enhanced affinity of ACE2^mod^-Fc could translate to improved biological functions, we conducted a pseudovirus neutralization assay. The neutralization profile of ACE2^mod^-Fc is apparently better compared to the profile of ACE2-Fc (Fig. 4a). There is more than a 10-fold improvement in both IC_50_ and IC_80_ values, comparing the two reagents. Anti SARS-CoV-2 immunoadhesin that binds to cell-surface displayed spike complexes might recruit beneficial immune factions via its Fc portion. We used flow cytometry to monitor the ability of ACE2-Fc and of ACE2^mod^-Fc to stain HEK293 cells that were transiently transfected express the SARS-CoV-2 spike complex (Fig. 4b). ACE2^mod^-Fc has an apparent higher capacity to recognize the spike complex compared to ACE2-Fc. Altogether, we created a superior ACE2-based immunoadhesin that displays an improved capacity to target SARS-CoV-2.

**Fig. 4.**
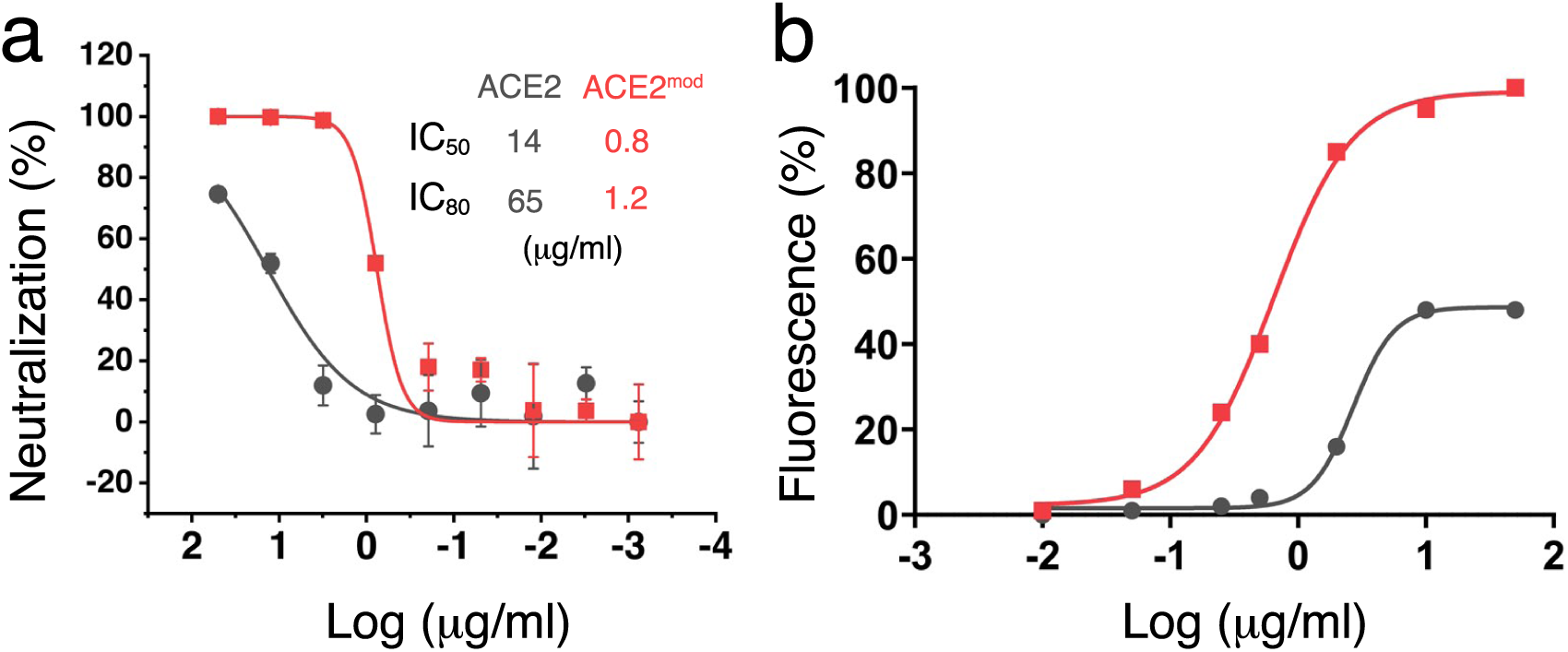
ACE2^mod^-Fc exhibits improved biological activity. **a**. Neutralization of pseudotyped viruses by ACE2-Fc (grey) and ACE2^mod^-Fc (red) is shown. The calculated IC_50_ and IC_80_ values are indicated. Error bars represent standard deviations. This experiment was repeated three times, and a representative graph is shown. **b**. Flagging of the SARS-CoV-2 spike complex on the cell-surface. The concentration-depended ability of ACE2-Fc (grey) and ACE2^mod^-Fc (red) to bind to surface-displayed SARS-CoV-2 spike complexes is analysed using flow cytometry. Normalized overall fluorescence signals for the gate-selected cells are shown. This experiment was repeated three times, and a representative graph is shown.

## Discussion

Immunotherapy makes a promising avenue to combat the COVID-19 pandemic. ACE2-based immunoadhesins were already described in the literature ^26,32^. However, reagents based on human-ACE2 have limited potency, and they can only compete in a stoichiometric fashion with the abundant natural ACE2 receptor on cells. An improved ACE2 binder, like the one we present here, is a more efficient competitor for binding to the SARS-CoV-2 RBD. Neutralization of the virus by blocking the receptor binding site is a mechanism that is shared by some anti-SARS-CoV-2 RBD mAbs that were recently isolated ^15-17^. Nevertheless, ACE2^mod^-Fc is likely to interact solely with the residues that make the ACE2 binding site, whereas the epitopes of anti-RBD mAbs are likely to only partially overlap with the ACE2-binidng surface, which may contribute for the emergence of escape mutations. Altogether we present a promising immunoadhesin that we now term Coronacept, which is a candidate immunotherapeutic agent for COVID-19.

## Methods

### Atomistic modeling

Orthologous sequences of ACE2 were collected by using a protein BLAST ^33^ search of the human-ACE2 sequence at GenBank and filtering the results to mammalian origin, and to sequences with greater than 80% identity to human-ACE2. Sequences were aligned using MUSCLE ^34^. Each sequence was threaded on the human-ACE2/Spike structure (PDB entry: 6VW1) and relaxed using sidechain packing and backbone, sidechain and rigid-body minimization subject to harmonic constraints on the Cα coordinates observed in the experimental structure. The ref2015 energy function was used in all calculations (xml and command line are provided in the Supplementary information). 100 models were generated for each sequence and the best scoring one was used for comparisons and analyses.

### Construction of expression vectors

Codon-optimized forms of human ACE2 binding region (amino-acids 19-615) and modified ACE2 genes were chemically synthesized (Genscript), and were subcloned upstream of a human Fc region (derived from IgG1) as previously described ^35^. Plasmid (pCMV3) encoding the Full-length Clone DNA of SARS-CoV-2 spike was purchased from Sino Biological, and subcloned to the same plasmid after removing 19AA from the C’-terminus (Δ19 S_covid-pCMV3, done by Lab of Yossi Shaul). Luciferase-pLenti6 and ΔR89 Ψ vectors for lentivirus production were a kind gift from the lab of Dr. Julia Adler (Prof. Shaul Lab, Weizmann Institute). The plasmid encoding the His-tagged SARS-CoV-2 Receptor Binding Domain (RBD) was a kind gift from Florian Krammer lab ^36^. Full-length human ACE2 was a kind gift from Hyeryun Choe ^22^ (Addgene plasmid #1786).

### Protein expression and purification

ACE-Fc fusion proteins and the SARS-CoV-2 RBD were expressed in suspension-HEK293F cells grown in FreeStyle media (Gibco). Transfections were done using 40 kDa polyethyleneimine (PEI-MAX) (Polysciences) at 1 mg of plasmid DNA per 1 L of culture at a cell density of 10^6^/ml. Media were collected six days post-transfection and supplemented with 0.02% (w/v) sodium azide and PMSF. S covid19 RBD was buffer exchanged to Phosphate Buffered Saline (PBS) using a tangential flow filtration system (Millipore), and captured using a HiTrap IMAC FF Ni^+2^ (GE Healthcare) affinity column followed by size exclusion chromatography purification with a Superdex 200 10/300 increased column (GE Healthcare). Fc-Fusion proteins were isolated using HiTrap protein-A (GE Healthcare) affinity columns.

### Surface Plasmon Resonance (SPR) measurements

Binding of SARS-CoV-2 RBD to ACE2 –Fc and ACE2^mod^–Fc fusion proteins were measured using a Biacore T200 instrument (GE Healthcare). Fusion proteins were first immobilized at a coupling density of ∽1000 response units (RU) on a series S sensor chip protein A (GE Healthcare) in PBS and 0.02% (w/v) sodium azide buffer. RBD was then injected at 0.16, 0.8, 4, 20, and 100 nM concentrations, at a flow rate of 60 μL/min. Single-cycle kinetics was performed for the binding assay. The sensor chip was regenerated using 10 mM glycine-HCl pH 1.5 buffer.

### Lentiviral particles production and Neutralization

Lentiviruses expressing S-Covid19 spikes were produced by transfecting HEK293T cells with Luciferase-pLenti6, Δ19 S_covid-pCMV3 and ΔR89 Ψ vectors at 1:1:1 ratio, using Lipofectamine 2000 (Thermo Fisher). Media containing Lentiviruses was collected at 48h post-transfection, centrifuged at 600g for 5min for clarifying from cells, and aliquots were frozen at -80°C.

For neutralization assays, HEK293T were transiently transfected with hACE2-pCDNA using Lipofectamine 2000. Following 18h post-transfection, cells were re-seeded on a poly-L-lysine pre-coated white, chimney 96-well plates (Greiner Bio-One). Cells were left to adhere for 8 h, followed by the addition of S-covid19 lentivirions, which were pre-incubated with 4-fold descending concentration series of either ACE2–Fc or ACE2^mod^–Fc. Luminescence from the activity of luciferase was measured 48 h post-infection using a TECAN infinite M200 pro plate reader after applying Bright-Glo reagent (Promega) on cells.

### Flow cytometry analyses

Flow cytometry was used to detect the binding between ACE2-Fc or ACE2^mod^-Fc to the spike complex of SARS-CoV-2 on cells. HEK293T cells were seeded on 10-cm plates and transfected 24 h later with 5 μg of Δ19 S_covid-pCMV3 using Lipofectamine 2000 (Invitrogen). Cells’ media were replaced 6 h later to full medium, i.e., DMEM (Biological Industries) supplemented with 1% Pen-Strep (v/v), 1% Glutamine (w/v), and 1% (v/v) MEM Non-essential amino acids. At 24 h post-transfection, cells were detached by pipetting and washed by centrifugation at 400xg for 5 min and re-suspension in PBS/0.5 % BSA solution. To prevent unspecific binding, blocking was performed by incubation in PBS/1 % BSA for 15 min. Cells were aliquoted and incubated with different concentrations of ACE2-Fc or ACE2^mod^-Fc diluted in PBS/0.5 % BSA: 50, 10, 2, 0.5, 0.25, 0.05, 0.01 μg/ml for 1 h, washed, and incubated with a 1:200 dilution of Alexa Fluor 488 donkey-anti-human secondary antibody for 30 min. Unstained cells and secondary antibody stained cells were used as negative controls. Analyses were performed using an LSR II flow cytometer (BD Biosciences). Curve fitting were performed using GraphPad Prism.

### ACE2 activity assay

ACE2 activity was evaluated using SensoLyte® 390 ACE2 Activity Assay Kit (ANASPEC; cat# 72086) according to the manufacturer’s protocol. 10 ng or 100 ng of ACE2-Fc and ACE2^mod^-Fc samples were compared blank control. Measurement of product formation (fluorogenic peptide cleavage) as a function of time was taken every 10 seconds.

## Acknowledgments

We are grateful to Julia Adler and Yosef Shaul for providing plasmids for the lentivirus system. Ron Diskin is the incumbent of the Tauro career development chair in biomedical research. This research was supported by the Ben B. and Joyce E. Eisenberg Foundation (R.D. & S.J.F.), the Ernst I Ascher Foundation, by a kind gift from Natan Sharansky (R.D.), and by Sam Switzer and family (S.J.F).

## Author contributions

R.D. conceived and oversaw the project; H.C.D produced and analyzed ACE2-Fc and its variants; J.W. and S.F. performed computer-based modeling and analysis; R.S. established the pseudotyped viral neutralization assay; M.S. performed enzymatic activity assay; M.E. and Y.M. performed FACS analyses; M.K., M.E., and A.S. assisted in molecular biology and tissue culture efforts; R.D. wrote the manuscript with comments from all other coauthors.

## Competing interests

The Weizmann Institute has filed for a patent for Coronacept.

