## Supplemental information for "Coronacept – a potent immunoadhesin against SARS-CoV-2"

This file contains:
Supplementary figure 1
Supplementary methods

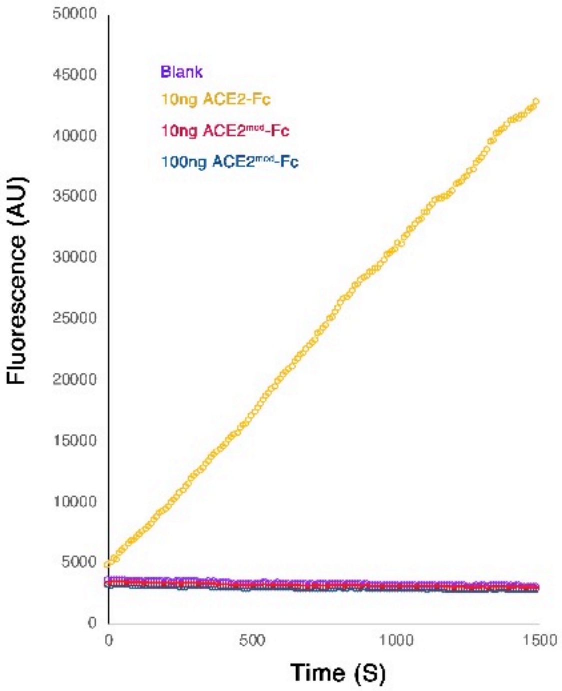


**Supplementary Figure 1 | Enzymatic activity of ACE2**. Enzymatic activity of ACE2-Fc and ACE2$^{mod}$-Fc was evaluated using an activity reporter assay. Activity of 10ng of ACE2-Fc, as well as 10ng and 100ng of ACE2$^{mod}$-Fc was measured and compered to blank control.

Supplementary methods

**Rosetta modeling of ACE2 orthologs** -
To model the structures of ACE2 orthologs the following command line was used: rosetta_scripts_executable -pdb_gz -s 6vw1_AE.pdb -parser:protocol refine_auto.xml - database Rosetta/main/database -parser:script_vars cst_value=0.4 -parser:script_vars cst_full_path=6vw1_AE_coord.cst -parser:script_vars scfxn=ref2015 -parser:script_vars soft_scfxn=soft_rep -use_input_sc -extrachi_cutoff 10 -ignore_unrecognized_res - chemical:exclude_patches LowerDNA  UpperDNA Cterm_amidation SpecialRotamer VirtualBB ShoveBB VirtualDNAPhosphate VirtualNTerm CTermConnect sc_orbitals pro_hydroxylated_case1 pro_hydroxylated_case2 ser_phosphorylated thr_phosphorylated tyr_phosphorylated tyr_sulfated lys_dimethylated lys_monomethylated  lys_trimethylated lys_acetylated glu_carboxylated cys_acetylated tyr_diiodinated N_acetylated C_methylamidated MethylatedProteinCterm -linmem_ig 10 -ignore_zero_occupancy false - load_PDB_components false

Where 6vw1_AE.pdb is a PDB file based on the PDB structure 6VW1 with chains A and E only, 6vw1_AE_coord.cst is a list of Ca atom coordinates from the above structure, and the refine_auto.xml script is:

```
<ROSETTASCRIPTS>
 <SCOREFXNS>
      <ScoreFunction name="ref_full" weights="%%scfxn%%">
      <Reweight scoretype="coordinate_constraint" weight="%%cst_value%%"/>
      </ScoreFunction>
      <ScoreFunction name="soft_rep_full" weights="%%soft_scfxn%%">
      <Reweight scoretype="coordinate_constraint" weight="%%cst_value%%"/>
      </ScoreFunction>
      <ScoreFunction name="ref_pure" weights="%%scfxn%%"/>
 </SCOREFXNS>
 <RESIDUE_SELECTORS>
      <Chain name="chain" chains="E"/>
 </RESIDUE_SELECTORS>
 <TASKOPERATIONS>
      <OperateOnResidueSubset name="rtr_E" selector="chain" >
      <RestrictToRepackingRLT/>
      </OperateOnResidueSubset>

      <InitializeFromCommandline name="init"/>
      <RestrictToRepacking name="rtr"/>
      <AlignedThread name="thread_to" query_name="%%query_name%%"
template_name="pdb6vw1.ent.A" alignment_file="/home/labs/fleishman/jonathaw/covid-
19/ace2-seq-analysis/short_aln.fasta" start_res="1"/>
 </TASKOPERATIONS>
 <MOVERS>
      <PackRotamersMover name="thread" scorefxn="soft_rep_full"
task_operations="init,thread_to,rtr_E"/>
```

```
	<Docking name="docking" score_high="ref_full" fullatom="1" local_refine="1"
jumps="1" task_operations="init,rtr,rtr_E"/>
	<PackRotamersMover name="soft_repack" scorefxn="soft_rep_full"
task_operations="init,rtr,rtr_E"/>
	<PackRotamersMover name="hard_repack" scorefxn="ref_full"
task_operations="init,rtr,rtr_E"/>
	<RotamerTrialsMinMover name="RTmin" scorefxn="ref_full"
task_operations="init,rtr,rtr_E"/>
	<MinMover name="soft_min" scorefxn="soft_rep_full" chi="1" bb="1" jump="0"/>
	<MinMover name="hard_min" scorefxn="ref_full" chi="1" bb="1" jump="0"/>
	<ConstraintSetMover name="add_CA_cst" cst_file="%%cst_full_path%%"/>
	<ParsedProtocol name="refinement_block"> #10 movers
	<Add mover_name="docking"/>
	<Add mover_name="soft_repack"/>
	<Add mover_name="soft_min"/>
	<Add mover_name="soft_repack"/>
	<Add mover_name="hard_min"/>
	<Add mover_name="hard_repack"/>
	<Add mover_name="hard_min"/>
	<Add mover_name="hard_repack"/>
	<Add mover_name="RTmin"/>
	<Add mover_name="RTmin"/>
<Add mover_name="hard_min"/>
	</ParsedProtocol>
	<LoopOver name="iter4" mover_name="refinement_block" iterations="4"/> #16
reacpk+min iterations total
  </MOVERS>
  <FILTERS>
	<ScoreType name="stability_score_full" scorefxn="ref_full"
score_type="total_score" confidence="0" threshold="0"/>
	<ScoreType name="stability_pure" scorefxn="ref_pure" score_type="total_score"
confidence="0" threshold="0"/>
	<Ddg name="a_ddg" scorefxn="ref_pure" chain_num="2" repeats="5"
extreme_value_removal="true" confidence="0" threshold="-5"/>
	<PackStat name="a_pack" confidence="0" threshold="0.3" chain="1"/>
	<BuriedUnsatHbonds2 name="a_unsat" scorefxn="ref_full" jump_number="1"
confidence="0"/>
	<ShapeComplementarity name="a_shape" confidence="0" jump="1"/>
	<BindingStrain name="a_bind" scorefxn="ref_full" jump="1" confidence="0"
threshold="5"/>
	<Sasa name="a_sasa" confidence="0" threshold="300"/>
	<Rmsd name="a_rmsd" confidence="0"/>
	<Time name="timer"/>
  </FILTERS>
  <PROTOCOLS>
	<Add filter_name="timer"/>
	<Add mover_name="add_CA_cst"/>
```

```
        <Add mover_name="thread"/>
        <Add mover_name="iter4"/>
        <Add filter_name="stability_score_full"/>
        <Add filter_name="stability_pure"/>
        <Add filter_name="a_rmsd"/>
        <Add filter_name="a_ddg"/>
        <Add filter_name="a_pack"/>
        <Add filter_name="a_unsat"/>
        <Add filter_name="a_shape"/>
        <Add filter_name="a_bind"/>
        <Add filter_name="a_sasa"/>
        <Add filter_name="timer"/>
  </PROTOCOLS>
  <OUTPUT scorefxn="ref_full"/>
</ROSETTASCRIPTS>
```
